# Chromogenic detection of telomere lengths in situ aids the identification of precancerous lesions in the prostate

**DOI:** 10.1101/2023.04.04.535575

**Authors:** Onur Ertunc, Erica Smearman, Qizhi Zheng, Jessica L. Hicks, Jacqueline A. Brosnan-Cashman, Tracy Jones, Carolina Gomes-Alexandre, Levent Trabzonlu, Alan K. Meeker, Angelo M. De Marzo, Christopher M. Heaphy

**Author notes:** **Correspondence to:** Christopher M. Heaphy, PhD, Boston University School of Medicine, 650 Albany Street, Room 444, Boston, MA 02118, or, Angelo M. De Marzo, MD, PhD, The Johns Hopkins University School of Medicine, CRB2 Room 144, 1550 Orleans Street, Baltimore, MD 21231. Department of Pathology, Suleyman Demirel University, Isparta, Turkey (Onur Ertunc). **Disclosures:** AMD is a paid consultant or advisor to Merck and Cepheid and has received research funding from Janssen and Myriad for unrelated work.

## Abstract

Telomeres are terminal chromosomal elements that are essential for the maintenance of genomic integrity. The measurement of telomere content provides useful diagnostic and prognostic information, and fluorescent methods have been developed for this purpose. However, fluorescent-based tissue assays are cumbersome for investigators to undertake, both in research and clinical settings. Here, a robust chromogenic *in situ* hybridization (CISH) approach was developed to visualize and quantify telomere content at single cell resolution in human prostate tissues, both frozen and formalin-fixed, paraffin-embedded (FFPE). This new assay (“Telo-CISH”) produces permanently stained slides that are viewable with a standard light microscope, thus avoiding the need for specialized equipment and storage. The assay is compatible with standard immunohistochemistry, thereby allowing simultaneous assessment of histomorphology, identification of specific cell types, and assessment of telomere status. In addition, Telo-CISH eliminates the problem of autofluorescent interference that frequently occurs with fluorescent-based methods. Using this new assay, we demonstrate successful application of Telo-CISH to help identify precancerous lesions in the prostate by the presence of markedly short telomeres specifically in the luminal epithelial cells. In summary, with fewer restrictions on the types of tissues that can be tested, and increased histologic information provided, the advantages presented by this novel chromogenic assay should extend the applicability of tissue-based telomere length assessment in research and clinical settings.

## INTRODUCTION

Located at chromosomal termini and bound by the six-member shelterin protein complex, telomeres are repetitive DNA elements (TTAGGGn) essential for maintaining genomic integrity (1-3). This telomere complex functions to inhibit exonucleolytic degradation, halt inappropriate homologous recombination, and prevent the chromosome ends from being recognized as double-strand breaks, thereby averting chromosomal fusions (4). In normal somatic cells, telomeres shorten with each cell division due to incomplete DNA replication on the lagging strand (5, 6). The cellular response to this telomere shortening depends on the status of DNA damage and cell cycle checkpoints (e.g., p53 and p16/RB). Normal cells possess surveillance systems that monitor telomere length and will induce either permanent cell cycle arrest (cellular senescence) or programmed cell death (apoptosis) in the event of telomeric DNA becoming critically shortened. These same cellular responses can also be triggered by an exquisitely sensitive DNA damage detection system capable of responding to telomere dysfunction-induced DNA damage, such as chromosome breakage-fusion-bridge cycles, thereby limiting cellular proliferative potential and preventing the accumulation of chromosomal instability (7). However, cells that have lost checkpoint controls may tolerate both critically short telomeres and the consequent DNA damage leading to chromosome breakage-fusion-bridge cycles resulting in the accumulation of genomic instability and thereby helping to foster malignant transformation (8-11). Consequently, to achieve the capacity for unlimited replication; a key cancer hallmark, the vast majority of cancers up-regulate the reverse transcriptase, telomerase, to stabilize telomeric ends (12), while a subset of cancers utilizes a telomerase-independent mechanism mediated by homology directed repair called Alternative Lengthening of Telomeres (ALT) (13, 14).

Abnormal telomere lengths are nearly ubiquitous in human pre-malignant lesions and cancers, most of which contain at least a subset of cells with markedly reduced telomere lengths (15-20). Several methods have been developed to analyze telomeric DNA in normal and cancerous cells (21-24). Previously, we and others developed a fluorescence *in situ* hybridization technique (TELI-FISH) allowing direct visualization and assessment of the lengths of telomeric DNA in formalin-fixed paraffin-embedded (FFPE) tissues (25-27). Advantages of this method over those utilizing extracted bulk genomic DNA (e.g., Southern blot-based and quantitative PCR methods) include the ability to assess telomere length at the individual cell level while preserving tissue spatial architecture, direct comparisons of telomere lengths in adjacent benign and cancer cells, simultaneous identification of cell types by immunofluorescence co-staining, and the use of minute archival tissue samples.

Since telomeric length abnormalities are a hallmark of human cancer and its precursors and are rarely found in benign counterparts, *in situ* telomeric length measurements are useful in a variety of clinical applications. Numerous studies have highlighted the translational potential of tissue-based telomere measurements for objectively identifying intraepithelial cancers, as well as for prognostication that might inform risk stratification for personalized therapeutic and surveillance strategies (28-31). While FISH provides excellent spatial resolution and the ability to obtain quantitative data, fluorescent microscopes are not part of the routine clinical workflow of surgical pathologists evaluating biopsy and resection specimens in real world settings. Furthermore, fluorescent assays require considerable expertise and substantial time to evaluate for quality control and detailed morphological analyses. An improved approach would allow for a more broadly accessible assessment of telomere lengths in both frozen and FFPE archival human tissues at a single cell resolution and can be routinely used in clinical laboratories with commercially available reagents. Chromogenic assays for telomere detection have previously been developed, although these assays have not been widely adapted, likely due to the fact that the assays do not rely on commercially available reagents and probes that are amenable to rapid adaptation and easily adopted for use on clinical autostainers (32, 33).

ACD RNAscope technology (Advanced Cell Diagnostics, Newark, CA) is a highly sensitive and specific *in situ* hybridization technique that can be used either in fluorescent or chromogenic assays on FFPE tissues (34). A variant of the technique, DNAscope, has been also used to detect viral DNA *in situ* (35, 36), as well as mitochondrial DNA (37). We have previously introduced telomere *in situ* hybridization using ACD DNAscope in two prior studies as a quality control measure (36, 37). In the present study, we developed and validated a chromogenic *in situ* hybridization assay using ACD RNAScope to allow for detection of telomeric DNA (“Telo-CISH”). Herein, we fully characterize this approach by showing its quantitative nature, verifying its applicability to the assessment of both cancerous and precursor lesions in the prostate. This new assay has several advantages over fluorescence-based methods, including the use of a standard light microscope, increased facile recognition of histomorphologic features that are difficult to discern using FISH probes, elimination of complications in interpretation due to tissue autofluorescence, and potential for clinical lab implementation.

## MATERIALS AND METHODS

### Cell culture

HeLa cells bearing a single copy of dox-inducible TPP-1 (38) were provided as a kind gift (J. Nandakumar). Cells were propagated in growth media containing DMEM, 10% FBS, 2 mM glutamax, 1 mM sodium pyruvate, and 1% penicillin/ streptomycin. Doxycycline was added to a final concentration of 200 ng/ml. Time points were taken from continuous growth in culture with the presence of Doxycyclin. In addition, three other cell lines (PC3, U-251, CHLA-200) representing a wide spectrum of telomere lengths were grown. PC3 and U-251 were obtained as part of the NCI-60 cancer cell line panel from the National Cancer Institute. CHLA-200 was obtained from the Children’s Oncology Group. PC3 and U-251 were grown in RPMI (Thermo Fisher Scientific) supplemented with 10% FBS and 1% penicillin/streptomycin. CHLA-200 was grown in IMDM (Thermo Fisher Scientific) supplemented with 20% FBS, 4 mM L-Glutamine, 1% penicillin/streptomycin, and 1X Insulin-Transferrin-Selenium. Media for CHLA-200 and U-251 also contained 1% Amphotericin B, 10 μg/mL Gentamicin, and 5 μg/mL Plasmocin (InvivoGen). Cells were submitted to the Genetic Resources Core Facility at The Johns Hopkins University School of Medicine for mycoplasma detection and cell line authentication by short tandem repeat (STR) profiling.

### Cell block preparation

FFPE cell blocks were generated using previously described protocols (36, 37). Briefly, cells were harvested, washed in phosphate buffered saline (PBS), then washed in 150 mL of 10% neutral buffered formalin (NBF). The resuspended cells were transferred to a 0.5 mL microcentrifuge tube with solidified 2% agarose prefilled in the tapered portion of the bottom, and spun in a swinging bucket centrifuge at 1000 rpm for 5 minutes. The resulting cell pellet was fixed in 10% NBF for 48 hours at room temperature by submerging the microcentrifuge tube into a 15 mL conical tube containing 10% NBF. After fixation, the microcentrifuge tubes were transferred to a new 15 mL conical tube with 10 mL PBS and stored at 4 degrees C prior to processing and embedding in paraffin blocks.

### Human tissue samples

A tissue microarray (TMA) was constructed from prostate tissues obtained from radical prostatectomies and normal control tissues through the Johns Hopkins Tissue Microarray Laboratory. Additional human prostate tissues (frozen and FFPE) were obtained from radical prostatectomies performed at The Johns Hopkins Hospital. This study was approved by the Johns Hopkins institutional internal review board.

### Telomere chromogenic in situ hybridization (Telo-CISH)

Chromogenic *in situ* hybridization (CISH) of telomeric DNA was performed manually following the instructions from the manufacturer (Advanced Cell Diagnostic; ACD) using the RNAscope Multiplex Fluorescent Reagent Kit v2 (Cat # 323100, ACD) with the following modifications. For FFPE tissues and cell line blocks, slides were baked at 60 degrees C for 30 minutes, deparaffinized in xylene 3 times for a total of 20 minutes, and in 100% ethanol 2 times for 4 minutes. Slides were then incubated in hydrogen peroxide solution for 10 minutes at room temperature (RT) and steamed in 1x RNAscope Target retrieval reagent at 100 degrees C for 18 minutes followed by protease plus digestion for 30 minutes at 40 degrees C. Slides were incubated in Hs-Telo-01 probe (Cat # 465451, ACD) at 1:2 dilution for 2 hours at 40 degrees C followed by the standard amplification steps as instructed by the manufacturer. After HRP C1, DAB solution was applied onto the slides for 10 min at RT. Of note, other telomere probes were tested; Hs-Telo (Cat # 465441, ACD), Hs-Telo-02 (Cat # 465461, ACD), and Hs-Telo-03 probe (Cat # 465471, ACD) gave strong nuclear signals consistent with telomeric DNA, but we did not pursue these probes further after initial testing. Thus, Hs-Telo-01 probe was utilized for all subsequent experiments.

For fresh frozen tissue sections, slides were prepared and pretreated following the instructions of the manufacturer (ACD) with the following modifications. Briefly, the tissue sections were fixed in pre-chilled 10% NBF at 4 degrees C for 15 minutes and dehydrated in a graded ethanol series. Next, the sections were pretreated in hydrogen peroxide for 10 minutes at RT, steamed in 1x RNAscope Target retrieval reagent at 100 degrees C for 5 minutes, incubated in protease IV for 10 minutes at room temperature, and then subjected to standard probe hybridization and signal amplification steps using RNAscope Multiplex Fluorescent Reagent Kit v2.

### Multiplex Telo-CISH/immunohistochemistry

In some cases, following CISH of telomeric DNA, multiplex IHC was performed. Immediately after performing Telo-CISH, we incubated the basal-specific cytokeratin primary antibody (Cat # ENZ-C34903; Enzo) at 1:50 dilution for 45 minutes at RT, followed by the PowerVision Poly-AP anti-mouse IgG secondary antibody (Cat # PV6110; Leica) for 30 min at RT. After primary and secondary antibody incubation, slides were rinsed with TBST. Next, ImmPACT Red (Cat # K5105; Vector Labs) was applied to the slides and incubated for 30 minutes at RT. Finally, the slides were counterstained with 50% Gill’s Hematoxylin for 30 seconds, quickly submersed in 100% ethanol and xylene, and then coverslipped with Cytoseal mounting medium.

### Fluorescence in situ hybridization (FISH)

Telomere lengths were assessed by fluorescent staining for telomeric DNA as previously described (25, 39). In brief, deparaffinized slides were hybridized with a Cy3-labeled peptide nucleic acid (PNA) probe (Cat # F1002, Panagene) complementary to the mammalian telomere repeat sequence ([N-terminus to C-terminus] CCCTAACCCTAACCCTAA). As a positive control for hybridization efficiency, an Alexa Fluor 488–labeled PNA probe (Cat # F3012, Panagene) with specificity for centromeric DNA repeats (CENP-B–binding sequence; ATTCGTTGGAAACGGGA) was included in the hybridization solution. Slides were counterstained with 4′,6-diamidino-2-phenylindole (Cat # D9542, Sigma-Aldrich) for labeling total nuclear DNA and mounted with Prolong anti-fade mounting medium (Cat # P36934, Life Technologies).

### Immunohistochemistry (IHC)

In some cases, IHC with either basal-specific cytokeratin (i.e. 903), PIN4, or MYC was performed on adjacent tissue sections as outlined. For 903, the Ventana Discovery Amp HQ kit and protocol which consists of a 32 minutes antigen retrieval step was used, followed by incubation for 40 minutes with 1:50 dilution of basal-specific anti-cytokeratin primary antibody (34BE12, Cat # ENZ-C34903, Enzo). After incubation, the HQ-MS step ran 12 minutes, and anti-HQ applied 12 minutes, then applied DAB. For PIN4, which consist of a cocktail mix of 903 (34BE12, Cat # ENZ-C34903, Enzo), P63 (Cat # CM163A, Biocare) and AMACR (Cat # Z2001L, Zeta), IHC was performed using the Discovery anti-HQ HRP kit (903/p63 cocktail) and the Discovery anti-HQ-NP kit (AMACR). The slides were steamed for 32 minutes in Cell Conditioning 1 (CC1) solution (Cat. # 950-500, Roche Diagnostics) for antigen retrieval. Then, the corresponding 903/p63 primary antibody mix was applied for 40 minutes at each at 1:50 and AMACR for 32 minutes at 1:50. DAB was applied and slides were coverslipped. For MYC, the Ventana Discovery Amp HQ kit and protocol which consists of a 48 minutes antigen retrieval step was used. The anti-c-Myc rabbit antibody (Y69, Cat # ab32072, Abcam) was incubated at 1:400 for 1 hour, then applied HQ-rabbit for 12 minutes, anti-HQ 12 minutes, followed by Amp 8/8 (2 steps) application. Finally, DAB was applied and slides were coverslipped.

### Microscopy

All chromogenic slides were scanned using a Roche-Ventana DP200 (Roche) or a Hamamatsu Nano-zoomer XR whole-slide scanner (Hamamatsu Photonics), and uploaded to Concentriq (Proscia) for whole-slide image viewing and micrograph documentation. For Telo-CISH images, HALO software (Indica Labs) and its ISH v2.2 and area quantification v1.0 algorithms were used to analyze the images. All fluorescent slides were scanned using the TissueFAXS Plus (Tissue Gnostics) automated microscopy workstation, which contains an 8-slide ultra-precise motorized stage and utilizes a Zeiss Z2 Axioimager microscope. For image analysis, we used the accompanying TissueQuest software module to analyze the fluorescent images with precise nuclear segmentation.

### Statistical analysis

Statistical analyses were performed using Stata for Windows software version 16 (StataCorp, College Station, TX). Pearson’s correlation coefficient (Pearson’s r) was used to investigate the relationship between Telo-CISH and quantification using Aperio based image analysis, as well as Telo-CISH and telomere FISH image analysis.

## RESULTS

### Development of a telomere-specific chromogenic in situ hybridization assay (Telo-CISH)

We sought to develop a quantitative telomere CISH assay that utilizes commercially available *in situ* hybridization probes and reagents compatible with both manual staining and automated protocols in a clinical pathology laboratory. We designed an Advanced Cell Diagnostics (ACD) probe set targeting the telomere repeats (Hs-Telo-01 probe). The application of this probe set for *in situ* hybridization displayed highly intense DAB signals with a punctate distribution within nuclei of prostate cells within FFPE human samples that were consistent with the known pattern and localization of telomeric DNA (**Figure 1A**). As a control during assay validation, pretreatment of slides with DNase I abolished the signal, confirming the specificity of the assay for DNA (**Figure 1B**).

**Figure 1.**
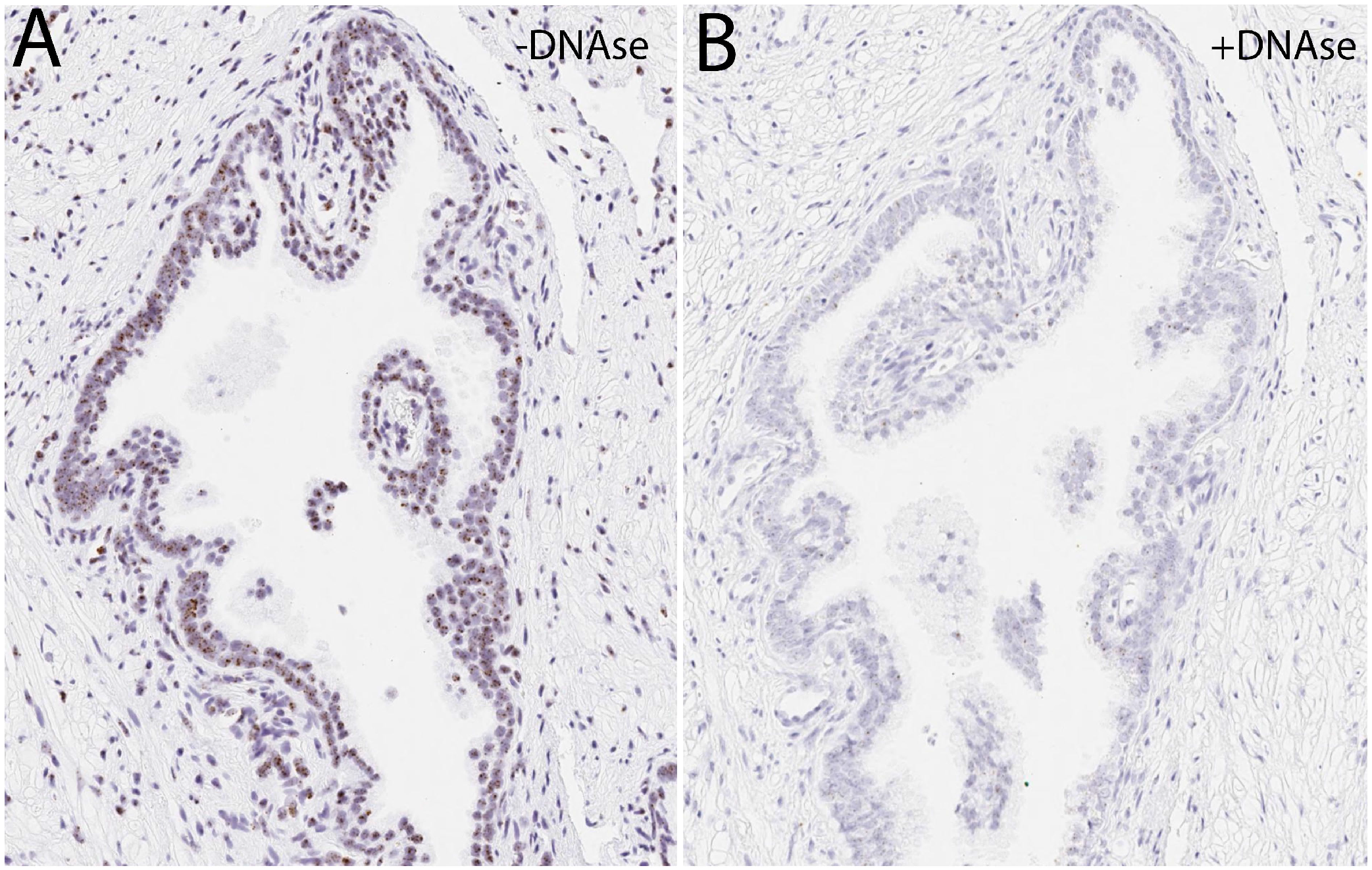
The specificity of the Telo-CISH assay verified by DNase pretreatment. Adjacent prostate cancer tissue slides hybridized with the human Hs-Telo-01 probe and visualized using the RNAscope 2.5 HD Assay-BROWN Detection Kit from ACD. Telo-CISH was performed on FFPE prostate tissues that were (**A**) untreated or (**B**) treated with DNase I. For both images, original magnification, ×200.

### Validation of Telo-CISH on human cell lines

To determine whether the Telo-CISH assay can provide a quantitative assessment of telomere content, we utilized a series of isogenic human cell lines with differing telomere lengths and subjected them to formalin fixation and paraffin embedding to simulate tissue fixation and processing (33, 36). As an initial approach, we employed isogenic HeLa cells with differing telomere lengths that are prepared by inducing the continuous expression of TPP-1, which results in gradually increasing telomere lengths over time (34). Cells were harvested cells after an increasing number of population doublings after TTP-1 induction and subjected to *in situ* hybridization with the Telo-CISH probe set. **Figure 2A** confirms that increasing amounts of telomeric DNA signal in the TPP-1 expressing HeLa cell lines with increasing telomere lengths can be visualized with Telo-CISH. **Figure 2B** shows the results of quantitative telomere FISH using our original assay with a PNA probe on these cells which, as expected, demonstrates a linear increase in telomere lengths in these isogenic cells with increasing population doublings. Next, we subjected the induced HeLa cells probed using the new Telo-CISH assay in Fig. 2A, as well as three other cell lines (PC3, U-251, CHLA-200) representing a wide spectrum of telomere lengths, to quantitative image analysis. This demonstrated a strong positive correlation between the Telo-CISH and telomere FISH results (**Figure 2C**) indicating that our novel chromogenic telomere CISH assay is quantitative and comparable to telomere FISH.

**Figure 2.**
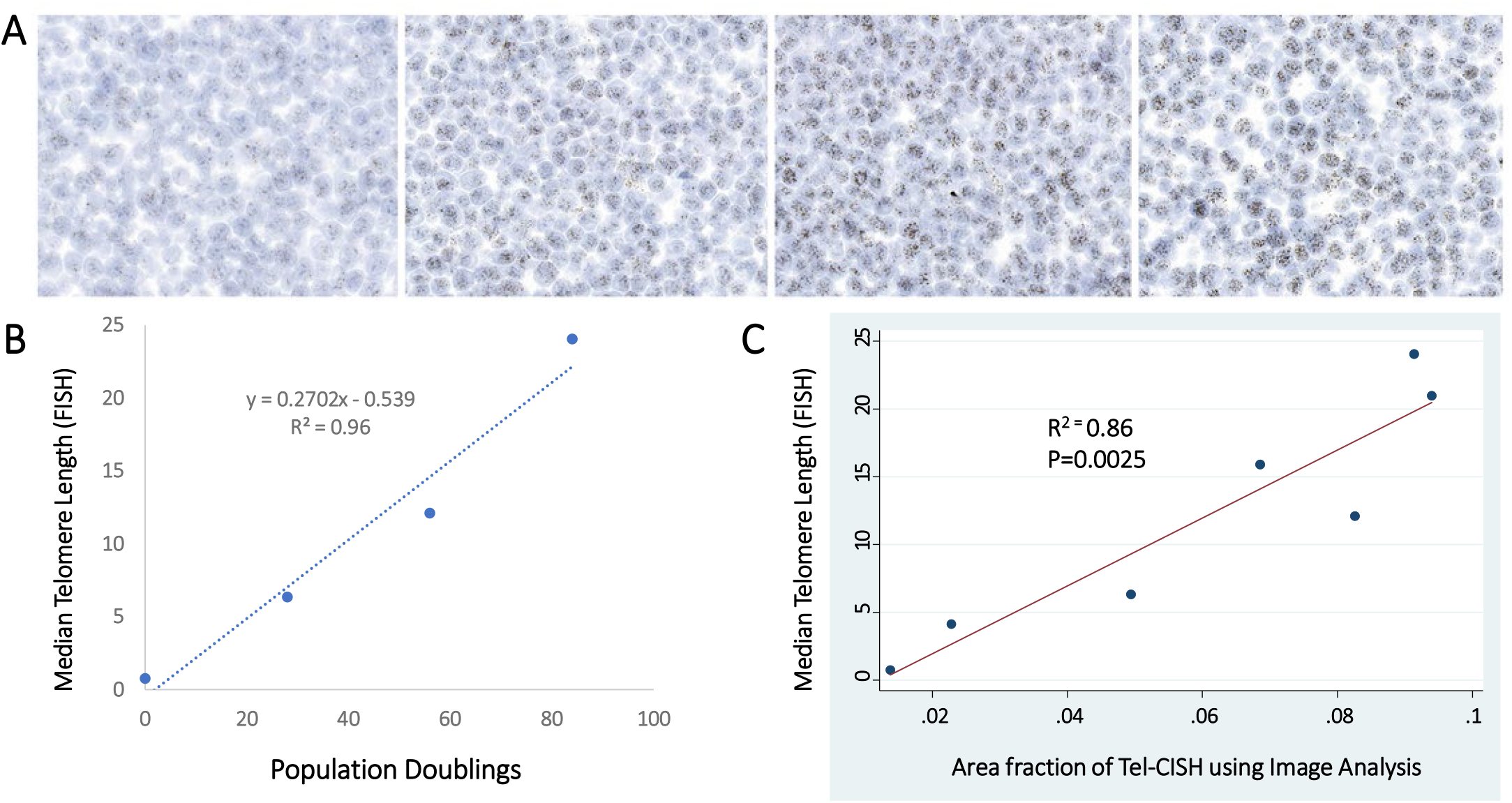
Quantitative assessment of the Telo-CISH assay. (**A**) HeLa cells bearing a single copy of dox-inducible TPP-1, known to increase telomere lengths, were continuously cultured. Cells were collected at different population doublings (PD0, PD28, PD56, PD84) for the construction of FFPE cell blocks. Telo-CISH was performed and a stepwise increase in telomeric signals was observed. (**B**) The FFPE cell blocks were also stained using TELI-FISH, and a positive correlation was observed between TELI-FISH-derived telomere lengths and population doublings (PD0, PD28, PD56, PD84). (**C**) Quantification of the Telo-CISH signals using Aperio based image analysis revealed a strong correlation with telomere FISH results in the four TPP-1 samples (PD0, PD28, PD56, PD84), as well as three additional cell lines representing a wide spectrum of telomere lengths (PC3, U-251, CHLA-200). Each dot corresponds to a different cell line. For all images, original magnification, ×400.

To determine whether the Telo-CISH assay is quantitative on human prostate tissues, we employed a small tissue microarray containing prostate cancer tissue and matched benign tissues for telomere FISH and Telo-CISH on adjacent slides. For both telomere FISH and Telo-CISH, quantitative image analyses revealed a strong positive correlation between the two assays, indicating that they are similarly quantitative in human tissues (**Figure 3A**). By visual inspection, both methods showed strong telomeric signals in the benign epithelium and stromal cells, with marked reductions in most cancer cells (**Figure 3B**).

**Figure 3.**
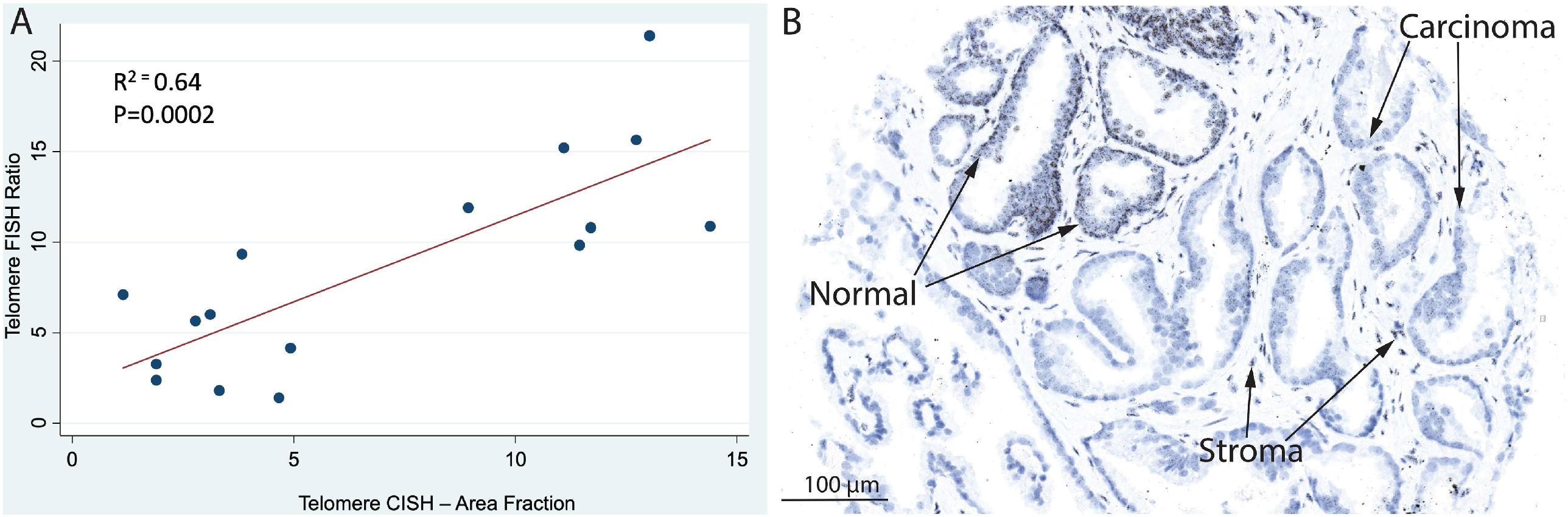
Direct comparison of the performance of the telomere FISH and Telo-CISH assays. Using a small tissue microarray consisting of prostate cancer and benign samples, (**A**) quantitative image analyses demonstrated a strong positive correlation between telomere FISH and Telo-CISH. (**B**) Example image of Telo-CISH in prostate cancer. Robust telomeric signals in the benign epithelial and stromal cells are present with the Telo-CISH assay, compared to marked reduction of telomeric signals in the cancer cells.

### Using Telo-CISH in frozen and archived human tissues

Since telomeres have been shown to be shortened in the vast majority of high-grade prostatic intraepithelial neoplasia (HGPIN) lesions, the presumptive precursor lesion to most prostatic adenocarcinomas, we sought to determine if Telo-CISH could reliably detect telomere shortening in archival standard FFPE sections from prostatectomy specimens. For this, we first demonstrated that Telo-CISH can be combined with immunolabeling for a basal-specific cytokeratin, which delineates normal prostatic basal cells, in a multiplex chromogenic assay. Using this assay, we show that cancer cells can be readily detectable by the presence of short telomeres as compared with adjacent normal luminal, basal, and stromal cells (**Figure 4**). Using this approach in a series of 12 prostatectomy specimens containing 19 HGPIN lesions, we found that all HGPIN lesions contained short telomeres. These prostatectomy specimens also contained 10 invasive adenocarcinomas (ranging from Gleason grade groups 1-5 and pathological stages T2N0-T3BN0). Consistent with previous observations, 9 of the 10 (90%) invasive adenocarcinomas showed markedly reduced telomere signals in the cancer cells compared with adjacent benign epithelium and stromal cells.

**Figure 4.**
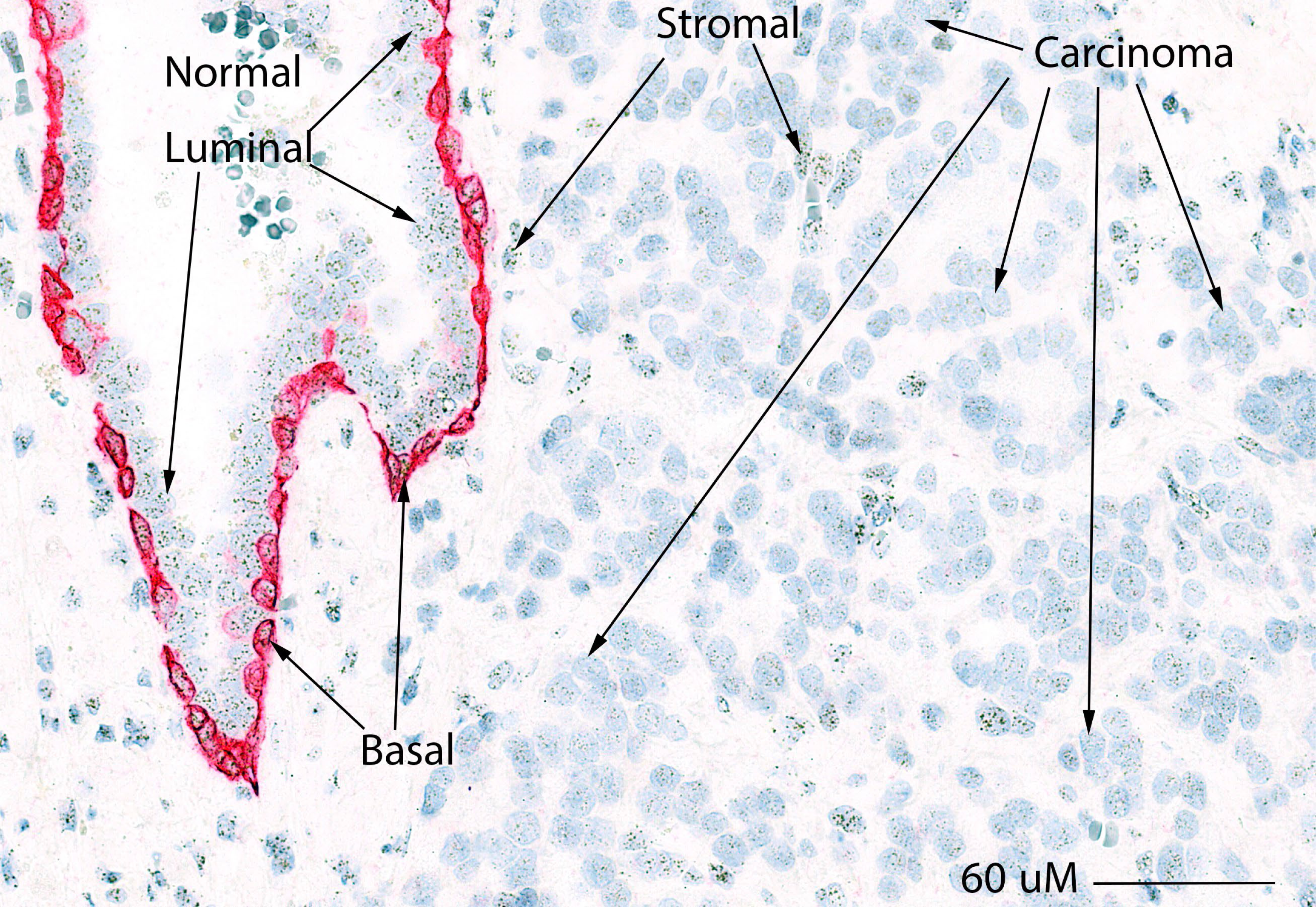
Telo-CISH is compatible for multiplexing with immunohistochemistry. Telo-CISH multiplex with immunohistochemistry for basal-specific cytokeratin which delineates the normal basal epithelial cells within the prostate (red chromogen), facilitates rapid scanning of whole slides to easily identify the presence of short telomeres in cancer cells. In this representative example, robust telomere signals are retained in the normal basal and luminal epithelial cells, as well as stromal cells.

Next, we sought to determine whether the combined Telo-CISH/IHC assay might have utility on frozen prostate tissue sections in order to aid in the diagnosis of HGPIN, which can be highly difficult due to the difficulty of discerning morphological details in frozen sections (37). We applied the Telo-CISH/IHC assay to a variety of fresh frozen human prostate tissues that contained a mixture of benign, precancerous (e.g., HGPIN), and cancerous regions. To validate the diagnosis of HGPIN independently, we employed IHC and *in situ* hybridization for MYC, which shows increased expression in luminal cells of HGPIN and can aid in its diagnosis (37). Using this approach, we evaluated 38 frozen section tissue blocks from 21 patients; the patient characteristics are outlined in **Table 1**. Gleason grade groups ranged from 3-5, pathological stages ranged from T2N0 to T3BN1, and patient ages ranged from 51 to 69. Of these men, 19 were White and 2 were Black. In total, we assessed 44 regions of HGPIN using a combination of H&E, PIN4 IHC (not shown), MYC IHC, MYC mRNA *in situ* hybridization (not shown), and all PIN lesions demonstrated telomere shortening compared to benign-appearing glands (**Figure 5**). Finally, in these same frozen tissue blocks, while there was some heterogeneity, 43 regions of cancer were identified and all (100%) showed marked telomere shortening in the majority of cancer cells compared to adjacent normal epithelium and stromal cells.

**Figure 5.**
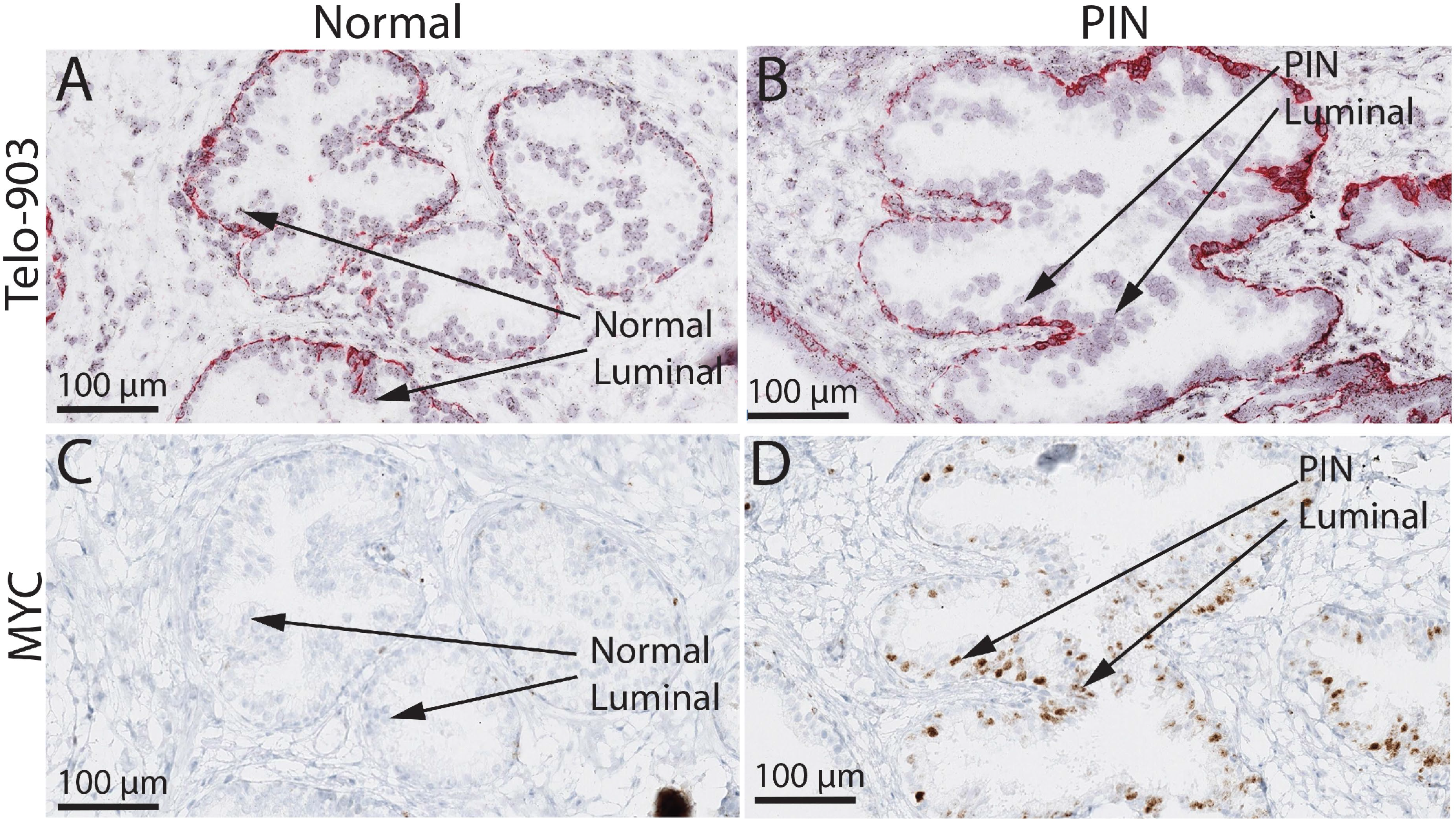
Tel-CISH facilitates the diagnosis of high-grade PIN lesions on frozen tissue sections. Tel-CISH multiplex with immunohistochemistry for basal-specific cytokeratin shows (**A**) robust telomere signals in the normal luminal epithelial cells of a benign prostatic gland, but (**B**) dramatically reduced telomeric signals in the luminal epithelial cells of a high-grade PIN lesion. To confirm the presence of high-grade PIN, MYC immunohistochemistry shows (**C**) lack of MYC protein in the normal luminal epithelial cells of a benign prostatic gland, but (**D**) increased MYC protein in the luminal epithelial cells of a high-grade PIN lesion.

**Table 1.** Telomere lengths in HGPIN and patient characteristics.

| **Case** | **Telomeres**<br>**(HGPIN)** | **Age** | **Race** | **Grade**<br>**Group** | **Pathological**<br>**Stage** |
| --- | --- | --- | --- | --- | --- |
| 1 | Short | 55 | Black | 2 | T2N0 |
| 2 | Short | 55 | White | 2 | T2N0 |
| 3 | Short | 61 | White | 2 | T2N0 |
| 4 | Short | 66 | White | 2 | T2N0 |
| 5 | Short | 66 | White | 2 | T2N0 |
| 6 | Short | 68 | White | 2 | T2N0 |
| 7 | Short | 56 | White | 2 | T3AN0 |
| 8 | Short | 58 | White | 2 | T3AN0 |
| 9 | Short | 59 | White | 2 | T3AN0 |
| 10 | Short | 57 | Black | 3 | T3AN0 |
| 11 | Short | 51 | White | 3 | T2N0 |
| 12 | Short | 67 | White | 3 | T3AN0 |
| 13 | Short | 67 | White | 3 | T3AN0 |
| 14 | Short | 56 | White | 3 | T3AN1 |
| 15 | Short | 66 | White | 3 | T3BN0 |
| 16 | Short | 54 | White | 5 | T3AN0 |
| 17 | Short | 66 | White | 5 | T3AN0 |
| 18 | Short | 69 | White | 5 | T3AN0 |
| 19 | Short | 55 | White | 5 | T3BN0 |
| 20 | Short | 63 | White | 5 | T3BN1 |
| 21 | Short | 59 | White | 5 | T3BN1 |

## DISCUSSION

Telomeres, the repetitive DNA elements located at chromosomal ends, shorten with each cell division, thereby playing an integral role in ensuring a finite cellular replicative capacity. In normal cells with intact cell cycle checkpoint proteins, progressive telomere shortening induces apoptosis or cellular senescence. However, abrogation of these checkpoints leads to aberrant cell proliferation and ultimately results in critically shortened telomeres. This, in turn, compromises chromosomal integrity by driving additional somatic copy number alterations, aneuploidy, and DNA rearrangements (40). Tissue-based measurements of telomeres have revealed that telomere shortening is prevalent in most human cancers, including prostate cancer (39). Additionally, telomere shortening is an early molecular event in cancer, occurring even in pre-malignant lesions (e.g. prostatic intraepithelial neoplasia, the presumed precursor lesion to prostatic adenocarcinoma) (15-18). Since short, dysfunctional telomeres promote genomic instability, the measurement of telomeres is useful in predicting disease progression in cancers such as prostate cancer (29, 30).

The direct assessment of telomere lengths in tissue samples, using fluorescently labeled probes has provided powerful tools to utilize in research applications (39). However, while valuable, these fluorescent-based methods have several drawbacks that have limited clinical applicability, including the need for costly and specialized equipment, loss of histologic detail, and a low signal to noise ratio due to the presence of tissue autofluorescence. Here, we developed a new method (Telo-CISH) to easily assess telomeres in either frozen or archival (FFPE) human specimens. Importantly, the new Telo-CISH assay utilizes commercially available reagents and produces permanently stained slides that are viewable with a standard light microscope, thereby avoiding the need for specialized microscopy equipment and special slide storage. We have demonstrated that this assay is compatible with standard hematoxylin and eosin staining, as well as immunohistochemical antibody stains, providing critical detailed histologic information. Importantly, Telo-CISH completely bypasses the significant problem of tissue autofluorescence. Finally, the assay is quantitative when performing image analysis and semi-quantitative visually such that a surgical pathologist can easily identify short telomeres in cancer cells or premalignant lesions compared to adjacent benign epithelial and stromal cells (**Figures 4 and 5**) Ongoing work is geared towards porting the assay to clinical automated strainers, so that that it can be readily employed in clinical laboratories using standard reagents. Since most epithelial precancerous lesions harbor short telomeres, this assay may prove useful in clinical trials focused on early disease detection and/or treatment.

In summary, we have developed a quantitative chromogenic *in situ* assay to assess telomeres (Telo-CISH) at the single cell level in frozen or archival human tissues. With increased histologic information, utilization of commercially available reagents, and need for only a standard bright-field microscope, these advantages will extend the applicability of tissue-based telomere length assessment in both research and clinical settings.

## ACKNOWLEDGEMENTS

This work is supported by NIH/National Cancer Institute (NCI) Specialized Programs of Research Excellence (SPORE) in Prostate Cancer grant P50CA58236 (AMD, AKM), NIH/NCI grant U01 CA196390 (AMD), NIH/NCI U54 CA274370 (AMD), U.S. Department of Defense Prostate Cancer Research Program (PCRP) grant W81XWH-18-2-0015 (AMD); The Johns Hopkins Sidney Kimmel Comprehensive Cancer Center Oncology Tissue Services Laboratory is supported by NIH/NCI grant P30 CA006973; The Patrick C. Walsh Prostate Cancer Research Fund at Johns Hopkins (AMD); Boston University School of Medicine Department of Medicine, Evans Medical Foundation (CMH). We wish to thank the staff at the Oncology Tissue Core and Microarray Core at Johns Hopkins Sidney Kimmel Comprehensive Cancer Center.

